# Real-time mass-resolved label-free single-molecule immunoassay

**DOI:** 10.64898/2026.02.17.706360

**Authors:** Carraugh C. Brouwer, Flaminia M. Muratori, Luke Melo, Luigi Anastasia, Carlo Pappone, Edward Grant

**Affiliations:** Department of Chemistry, University of British Columbia, Vancouver, BC V6T 1Z1 Canada; Biomedical and Materials Engineering, Politecnico di Milano, 20133 Italy; Institute for Molecular and Translational Cardiology (IMTC), IRCCS Policlinico San Donato, Milan, 20097 Italy; School of Medicine, University Vita- Salute San Raffaele, Milan, 20132 Italy

**Author notes:** Correspondence to Edward Grant Corresponding author.

## Abstract

Protein-protein interactions govern the biomolecular logic of immunity, signalling, and disease. Elementary rates of association, dissociation, and inhibition underpin our understanding of life processes and remain a limiting factor at the frontier of therapeutic discovery. Yet existing assays infer these processes indirectly through ensemble-averaged signals or molecular labels that obscure native dynamics. Here, we introduce a label-free, real-time, mass-resolved single-molecule immunoassay based on interferometric scattering microscopy (iSCAT) that directly observes individual protein-protein binding events in complex biological samples. By detecting light scattered from single proteins as they bind to an antibody-functionalized surface, we resolve discrete antibody-antigen interactions with single-molecule sensitivity and molecular-weight discrimination. Using IgM as a model system, we demonstrate real-time detection of individual binding events, with measured association rates that scale linearly with concentration over three orders of magnitude. Direct counts of IgM binding events in human serum yield quantitative concentrations that agree with bulk measurements obtained by enzyme-linked immunosorbent assay (ELISA). This bioaffinity iSCAT platform unifies molecular specificity, label-free detection, real-time kinetics, and mass-resolved single-molecule sensitivity, enabling direct access to protein-protein recognition processes and establishing a general framework for quantitative, single-molecule immunoassays.

---

Protein-protein interactions lie at the heart of nearly every biological process, governing how cells sense, respond, repair, and communicate. Their dysregulation drives a wide range of diseases, from autoimmunity to cancer and neurodegeneration. Resolving these interactions with molecular specificity and temporal precision remains a major unmet need in biophysics and bioanalytics with significant implications for disease diagnosis, therapeutic drug monitoring, and drug discovery (*1–4*). Immunoassays exploit highly specific antigen-antibody interactions to isolate and quantify target antigens in complex biological samples. Recent improvements include the development of label-free approaches (*5*), as well as methods with single-molecule sensitivity (*6–10*) and real-time methods offering the capacity to determine kinetic binding constants (*11, 12*). Yet, despite decades of innovation in immunochemistry, no immunoassay affords label-free, real-time, direct, mass-resolved single-molecule detection simultaneously.

In a novel use of interferometric scattering microscopy (iSCAT), we overcome this long-standing limitation by enabling real-time, label-free, mass-resolved detection of discrete antibody-antigen binding events on an antibody-derivatized and blocked coverslip. This approach transforms a routine iSCAT measurement into a true single-molecule immunoassay that quantifies target proteins in complex fluids and distinguishes molecular species by mass, without labels, amplification, or ensemble averaging. This label-free optical detection method avoids variability, steric perturbation, and kinetic distortion introduced by fluorescent or enzymatic tags, enabling direct measurement of native binding events. Single-molecule detection resolves binding heterogeneity, stoichiometry, and oligomeric state, enabling insights inaccessible to bulk assays. Real-time monitoring captures the dynamic sequence of adsorption and recognition, opening a path to genuine single-molecule kinetics on immuno-functionalized surfaces.

Conventional immunoassays share similar biorecognition principles but employ various detection platforms (*2*). Typically, antibodies bind to a surface, and a blocking agent occupies the remaining binding sites. Upon sample introduction, target molecules bind to the immobilized antibodies. Often, detection requires a secondary antibody with a molecular label. In enzyme-linked immunosorbent assay (ELISA), enzymatic labels catalyze a colour-changing reaction that is proportional to the analyte concentration, yielding a bulk, end-point measurement. Digital ELISA gains single-molecule resolution by confining analyte molecules to microbeads or microdroplets (*13*). Surface plasmon resonance detects analyte without the use of labels, producing a bulk signal that arises from a change in refractive index (*14*). Localized surface plasmon resonance can register single-molecule binding by micromanipulating analytes and secondary probes tethered to nanoparticles (*7*).

By contrast, label-free, single-molecule iSCAT directly detects light scattered by individual macromolecules alone as they land on the surface of a coverslip. The interferometric signal, defined by the scattered field combined with a specularly reflected reference field, *E*_*r*_, determines the iSCAT contrast 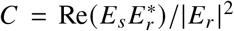 of a protein. The amplitude of scattered light, *E*_*s*_, varies with polarizability, which scales linearly with molecular mass, enabling quantitative optical mass measurements of single biomolecules (*15–20*)

Here, we demonstrate a direct, label-free, mass-resolved, real-time immunoassay in which an iSCAT measurement detects binding events on an antibody-derivatized surface. The present work proves the principle with a single-molecule immunoassay of an immunoglobulin M (IgM) model system. Experimental measurements detect and count individual IgM molecules landing on the capture surface with linear concentration dependence, distinguish IgM from IgA in mixed samples by their mass-dependent contrast signatures, and quantify endogenous IgM in human serum with accuracy comparable to ELISA, while remaining insensitive to non-target serum proteins.

Taken together, these findings demonstrate that bioaffinity iSCAT can serve as a true single-molecule immunoassay platform, delivering label-free molecular identification, real-time binding trajectories, and single-protein quantification directly in complex biological samples. By enabling direct observation of discrete, mass-resolved immuno-recognition events, this approach opens a new experimental window for probing protein-protein interactions, accelerating the development of therapeutic strategies targeting molecular recognition pathways in health and disease.

## Results

### Simultaneous measurement of IgA and IgM

Our custom-built iSCAT microscope images diffraction-limited scattering point-spread function signals formed in real time by individual protein molecules as they bind to a derivatized coverslip surface. Our particle tracking algorithm analyzes video records of such signals to log the location, adsorption time, and contrast of each binding event. The contrast distribution observed for a given protein forms a histogram centred at a value proportional to its molecular weight.

Figure 1 demonstrates this principle. For reference, Panel B shows contrast histograms observed in separate 30-second measurements of landings from 2 nM IgA and 1nM IgM solutions on coverslips bearing conventional poly-D-lysine (PDL) thin films at a pH of 7.4. The dominant pentameric form of IgM has a molecular mass of 970 kDa. This dimerizes to form a small concentration of decamer. The contrast histogram shows a dominant peak centred at 0.0033 and a secondary feature at 0.0066. The dimeric form of IgA has a molecular mass of 385 kDa. The IgA solution yields a single particle contrast distribution that peaks at 0.0013.

**Figure 1:**
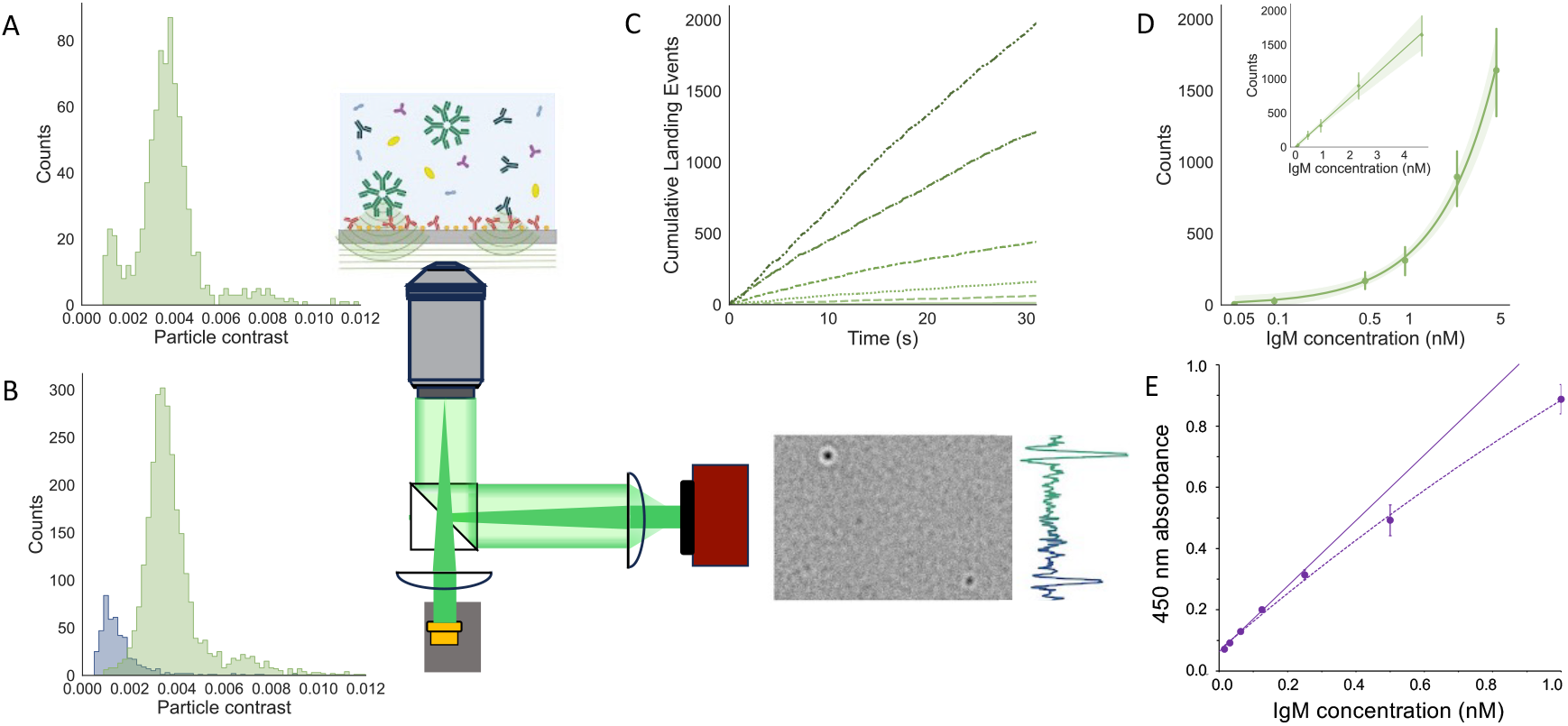
Immunoassay of IgM and IgM - IgA mixtures. A) Contrast histogram of landings from a solution containing 1 nM IgM and 2 nM IgA on an immunoassay-functionalized coverslip. B) Contrast histograms for 1 nM IgM (green) and 2 nM IgA (blue) on separate PDL-coated coverslips. C) Cumulative IgM landing events on immunoassay coverslips over time for IgM pentamer concentrations 0.046, 0.093, 0.46, 0.93, 2.3, and 4.6 nM in PBS. D) Standard curve correlating the IgM concentration with mass-resolved immunoassay landing rate on a log scale with a linear inset. E) Conventional ELISA analysis of an IgM concentration series in PBS. The schematic diagram depicts a common-path interferometric measurement in which a derivatized coverslip selectively binds single IgM pentamers and IgA dimers, which appear in real time as transient point-spread functions.

The mass discrimination of iSCAT enables simultaneous IgM and IgA immunoassay on a coverslip derivatized with a polyclonal anti-Ig antibody. Panel A shows the contrast histograms observed in a 30-second measurement of a solution containing 2 nM IgA dimer and 1nM IgM pentamer on an immunoassay-functionalized coverslip. Here, we see the same pattern of landings. Three evident peaks with contrast values centred at 0.0013, 0.0033, and 0.0066 match the individual contrast values observed in separate analyses of IgA and IgM solutions on non-discriminating PDL coverslips.

### Quantification of IgM concentration by particle counting

To test whether the observed iSCAT landing rate accurately gauges protein concentration in solution, we counted the appearance of single-molecule signals as a function of time for a wide range of IgM concentrations in PBS. Figure 1C displays the accumulation rate of IgM molecules on immunoassay functionalized coverslips for different IgM concentrations in PBS. We find that binding events increase linearly with IgM concentration over the full range from 0.046 nM to 4.6 nM. Individual observations of landing rate as a function of time show no plateau indicative of a depletion layer (*21*) within the first 30 s of measurement. Figure S2 in the Supplementary Materials shows single frames taken from video records of single-molecule IgM – anti-IgM binding events over this range of concentrations in PBS. Here, we also provide a link to corresponding 200-frame videos.

Counts of total numbers of bound molecules observed in 30-second measurements at various IgM concentrations in PBS form the basis of a standard curve. Figure 1D shows the concentration-dependent response in the total number of observed IgM binding events obtained by following the immunoassay procedure using known concentrations of IgM in PBS. We see a strong positive linear correlation between IgM concentration and IgM particle counts. The equation of the trendline is *y* = 360*x* − 0.36. Notably, this relationship spans three orders of magnitude. The standard error increases with IgM concentration. Note the intercept near zero and high correlation coefficient (*R*^2^ = 0.997), affirming the quantitative accuracy of this assay.

Compare this with the results of the conventional ELISA-based standard-addition curve plotted in Figure 1E. Here, note that the 450 nm absorbance response measuring IgM concentration exhibits a pronounced curvature above 0.2 nM. Notice as well that the standard-addition response intercepts the *y*-axis at a point that denotes a measured, nonzero response for a PBS blank.

### Binding specificity

Several negative control experiments serve to confirm the binding specificity of the iSCAT-based IgM immunoassay. Negative control experiments testing for signal using solutions of PBS buffer alone establish that coverslips derivatized with IgM antibody and blocked with casen yield a negligible background signal. In 30-second observations, PBS buffer alone produced an average of 0.3 landing events on casein-blocked control coverslips and 1.4 on derivatized and blocked coverslips. Other control experiments test the effectiveness of casein in blocking protein landing events in the absence of antibody derivatization. Buffered IgM solutions, normal serum, and depleted serum samples, all with high solute concentrations, exhibited negligible landings on casein-blocked coverslips. A 5 nM IgM sample produced 99.5% fewer landings on a casein-blocked coverslip than on a poly-D-lysine-derivatized coverslip. In a 30-second measurement, normal and depleted serum samples exhibited fewer than 100 and 50 landings, respectively, over a 30-second period. Finally, in a test for nonspecific binding to the capture antibody, a 5 nM ferritin solution yielded an average of fewer than 2 landings over a 30-second measurement. Supplementary Materials Table S1 summarizes the results of control experiments.

### Immunoassay selectivity for IgM and IgA in human serum

Tests using diluted human serum samples on different functionalized surfaces: poly-D-lysine (PDL) and anti-Ig with casein blocking, verify the specificity of the immunoassay when presented with a complex sample matrix. These experiments examined both normal serum and serum depleted of IgM, IgA, and IgG. Figure 2 shows the contrast values of observed landings for serum.

**Figure 2:**
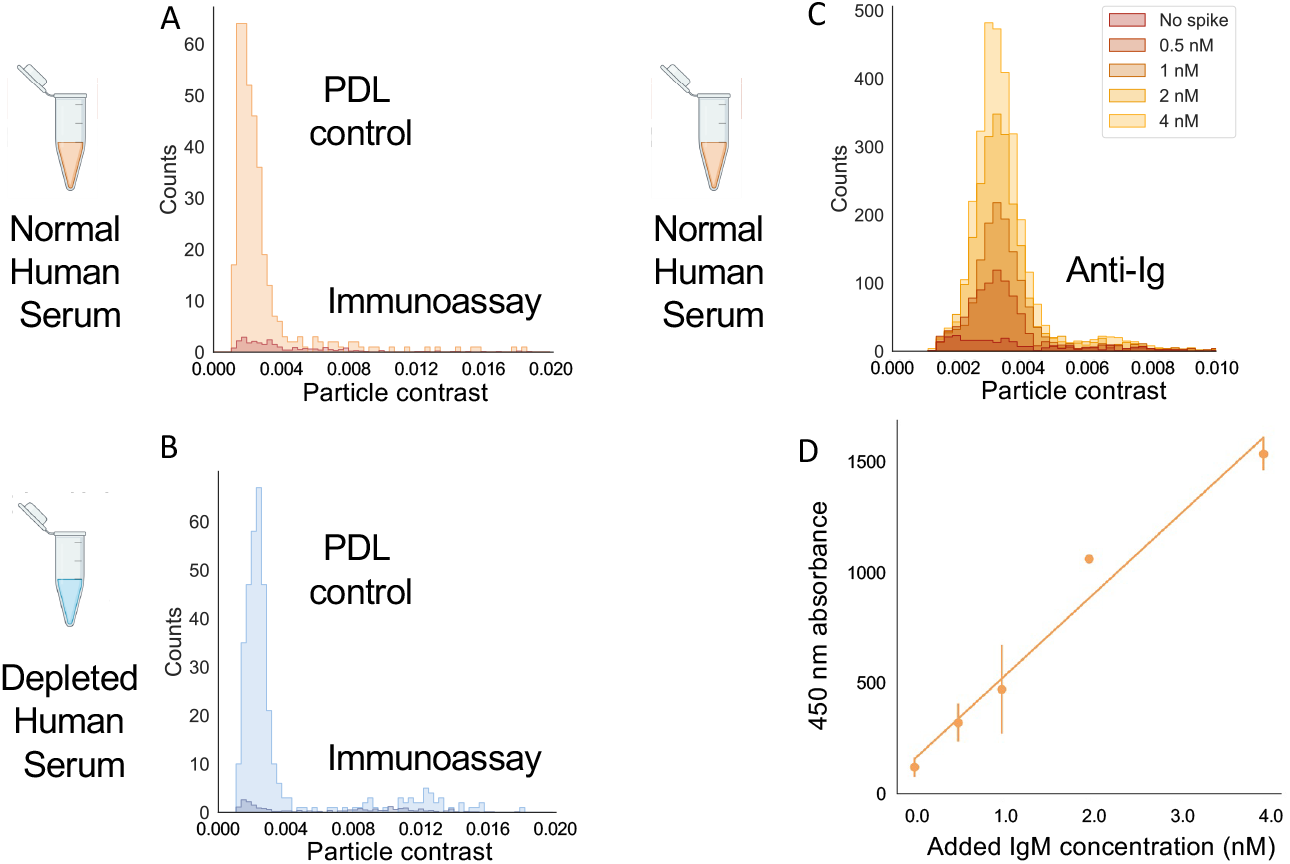
Immunoassay selectivity in human serum. Contrast histograms comparing numbers of landings observed in 30 seconds from human serum samples, diluted 1:5,000, on poly-D-lysine (light) and equivalent measurements on prepared immunoassay coverslips (dark). A) Normal serum B) IgM/IgA/IgG depleted serum. C) Contrast histogram showing unspiked serum and a series of IgM spikes. D) Linear regression relating the amount of IgM spike to the observed counts of IgM. The equation of the line is *y* = 371*x* + 121 with *R*^2^ = 0.967.

Normal serum produces numerous landings on PDL, forming a broad contrast distribution centered around 0.002. Few landings occur with higher contrasts. On an antibody-derivatized coverslip, normal serum forms iSCAT contrast histograms with peaks between 0.001 and 0.002 and between 0.003 and 0.004. Depleted serum shows a comparable contrast distribution on PDL. In contrast to the normal serum, depleted serum produces few landings with contrasts between 0.003 and 0.004 on an antibody-derivatized coverslip.

To ascertain the robustness of the iSCAT single-molecule immunoassay in a complex sample matrix, we added IgM spikes of known concentration to human serum. IgM spikes in serum increased the height of the peak centred in contrast histograms at 0.0033 (Figure 2C). The slope of the linear fit to the particle counts from varying added concentrations of IgM in serum (Figure 2D) is 371 particles/nM added IgM, which compares well with the slope of the calibration line obtained for IgM in PBS buffer (360 particles/nM IgM). The standard-addition intercept indicates a native IgM concentration of 163 mg/dL in the undiluted serum. A conventional sandwich enzyme-linked immunosorbent assay (ELISA) applied to the same human serum sample yields a comparable IgM concentration of 167 mg/dL ± 3 mg/dL.

## Discussion

This work introduces a novel, real-time, label-free immunoassay with single-molecule sensitivity. A custom-built interferometric scattering microscope directly detects individual antigen binding events. We demonstrate this assay on an IgM and anti-Ig model system. The analyte can be quantified based on the number of analyte adsorption events on an antibody-coated surface. Measured particle counts for nanomolar concentrations of IgM using the immunoassay procedure yield a standard curve (Figure 1D) with strong linearity (*R*2 = 0.997) over three orders of magnitude. Rates of landing are linear for 30 s following sample injection (Figure 1C), indicating that the number of available binding sites is not substantially reduced nor is the local concentration of IgM near the coverslip significantly depleted during that time. (*21*)

By comparison, the ELISA offers a much less ideal analytical response. The 450 nm absorbance signal response becomes visibly nonlinear for IgM concentrations above 0.2 nM. In addition, the calibration of IgM in blank PBS yields a nonzero *y*-intercept. This signal, proportional to enzyme turnover rather than direct count, can arise from an absorbance offset owing to nonspecific absorption or incomplete removal of enzyme-linked detection reagents during washing or optical baseline contributions inadequately removed by blank subtraction (*22, 23*).

The demonstrated linear dynamic range for iSCAT single-molecule detection, from 0.046 nM to 4.6 nM, is 10x higher than that measured with a commercial IgM ELISA kit. A low landing rate within the field of view sets the current lower measurement limit. Expanding the illumination field of view or extending the landing window duration would significantly improve the detection limit. Too many particles landing in close succession at higher concentrations can lead to issues with particle detection because point-spread functions may overlap, making individual particles indistinguishable (*24*). A higher frame rate or smaller ratiometric binning size could increase the upper detection limit by temporally separating particles in the ratiometric video. However, a higher frame rate makes smaller proteins harder to detect because the signal-to-noise ratio decreases with shorter exposure times (*19*).

This real-time mass-resolved label-free single-molecule immunoassay is specific to the target protein. In control experiments with high IgM concentrations, casein effectively blocks empty binding sites on the microscope coverslip. The average number of landings recorded in these negative controls is two orders of magnitude lower than that in positive controls. Control experiments testing non-specific protein binding to anti-Ig using ferritin further support assay specificity because no ferritin landings are recorded. Experiments on poly-D-lysine-functionalized coverslips show that serum biomolecules bind rapidly to PDL. PDL has a strong positive charge at neutral pH, and electrostatically adsorbs negatively charged species from solution. Most proteins have regions of negative charge, so PDL is considered a nonspecific attractor. The broadness of the low-contrast peak from serum molecules bound to PDL (Figure 2A) indicates the presence of a diversity of serum biomolecules with different masses. Comparatively few serum biomolecules bind to immunoassay-functionalized coverslips. The contrast distribution for the normal serum on the immunoassay surface has two main peaks around 0.002 and 0.004, corresponding to measured average contrasts of IgA and IgM (Figure 2B). In the depleted serum, where IgM has been removed, the peak around is no longer present. This observation suggests that IgM was successfully removed, and no other species of similar mass has affinity for the immunoassay surface.

IgM particle count rates increase with increasing IgM concentration equivalently in diluted serum samples and pure PBS buffer (cf. Figures 1D and 2D). Thus, other serum biomolecules have minimal impact on the immunoassay signal, enabling accurate IgM measurement in complex sample matrices. The *y*-intercept of the iSCAT standard addition curve (Figure 2D) determines the IgM concentration in undiluted normal serum to be 163 mg/dL. This value accords with typical IgM serum concentrations (*25*). A commercial ELISA determination of IgM in the present human serum sample yields a concentration of 167 mg/dL ± 3 mg/dL, further validating the accuracy of the iSCAT immunoassay.

Label-free detection provides mass resolution, enabling the detection of multiple analytes. A mass calibration curve relates iSCAT contrast to protein mass (Figure S2). The strong linear relationship between mass and contrast (*R*^2^ = 0.994) differentiates proteins without requiring labels. We demonstrate simultaneous detection of two different targets (IgM and IgA) of the iSCAT immunoassay using the anti-Ig antibody, which is known to bind to both IgM and IgA (Figure 2). The ability to identify IgA and IgM in serum without additional labelling steps reduces assay complexity, lowers costs, and preserves the native state of the proteins, avoiding potential alterations caused by tags or conjugates. Moreover, mass information from single measurements provides insight into oligomeric states of detected proteins. The reach of such experiments depends on the optical mass resolution of the iSCAT instrument (*18*). We expect that hardware and data-processing improvements underway will increase the mass resolution of the current instrument by a factor of five, and substantially lower its detection threshold.

The ability of immunoassays to detect specific proteins in complex samples makes them invaluable tools for biomolecular research and clinical diagnostics. However, the reliance of common methods on fluorescent labels and bulk measurements makes assays complex and time-consuming, and limits the scope of information that can be obtained from them. The addition of simultaneous label-free, real-time, single-molecule capability enhances the versatility of an immunoassay, offering direct, single-molecule, immunoselective measurements in real time.

Sample preparation requires less time and demands fewer resources than conventional methods, such as ELISA, because immunosorbent iSCAT detects target proteins without indicators or fluorescent tags. Assays that take no more than three hours in total provide instantaneous landing rates that indicate the concentrations of target analytes in solution. The relationship between protein mass and the scattered-light contrast of label-free proteins enables simultaneous detection of multiple immunologically active analytes in a single assay.

The immunosorbent landscape offers additional advantages. An iSCAT measurement resolves the *x, y* coordinates of every landing. The unbinding of a protein yields an easily recognized inverted contrast signal. The distribution of dwell times in a record of spatially resolved binding and unbinding events provides a direct, single-molecule readout of the associated *k*_off_. The selectivity of immunospecific binding also enables iSCAT measurements to determine protein-protein *k*_*d*_ values across regimes where high ligand concentrations would otherwise swamp the assay of free target.

## Methods

### Sample materials

Goat anti-Human IgG, IgM, IgA (H+L) Secondary Antibody (anti-Ig capture antibody, Cat: 31128), Human Immunoglobulin M Isotype Control (IgM, Cat: 31146), Human Immunoglobulin A Isotype Control (IgA, Cat: 31148), and Normal Human Serum (Cat: 31876) were purchased from ThermoFisher Scientific. Serum minus IgA/IgM/IgG human (depleted serum, Cat: S5393) was purchased from Millipore Sigma. Ferritin protein, in stock solution of 20 mg/mL, was provided by the Lallous laboratory at the Vancouver Prostate Centre. 1% casein solution (Cat: 37528) and phosphate-buffered saline (PBS, Cat: AM9625) solution were obtained from ThermoFisher Scientific. PBS and casein solution were filtered with a 0.22 *μ*m Millex-GP syringe filter (Millipore Sigma, Cat: SLGP033RS). Casein was centrifuged for 12 mins at 14,000 × g with 30 kDa Amicon Ultra-0.5 mL centrifugal filters (Millipore Sigma, Cat: UFC503008). Stock IgM (4.5 mg/mL) was diluted in PBS to concentrations between 0.046 nM and 4.64 nM. Anti-Ig capture antibody was diluted in PBS to 2 µM concentration. Ferritin was diluted in PBS to 5 nM and 7.6 nM. Human serum samples were centrifuged at 15,000 × g for 10 mins and diluted 5000-fold in PBS.

### Immunoassay surface preparation

A 4-well silicon gasket (Grace Bio-labs, Cat: GBL103250-10EA) is placed on a poly-D-lysine (PDL) coated glass coverslip (Neuvitro, Cat: GG-25-15-PDL). 10 *μ*L of anti-Ig capture antibody was pipetted in each well and incubated for 1 h at room temperature. Gaskets were covered with clean glass coverslips to avoid sample evaporation during the incubations. Three washes per sample with PBS removed unbound antibodies. 10 *μ*L of 1% casein solution was pipetted into each gasket and incubated for 1 h at room temperature. Six PBS washes per sample removed unbound casein.

### iSCAT sample measurement

Prepared coverslips are placed on the sample stage, with one silicon gasket centred above the microscope objective. 10 *μ*L of sample solution is added, and a 1 min video is recorded with 2 ms exposure time, a frame rate of 500 fps, and a field of view of 8.4 x 8.4 µm^2^.

### Ratiometric image processing

To process thousands of protein landings per minute, we employ ratiometric processing (*15– 17, 19*), which reveals dynamic changes in the interferometric image of light scattered by the coverslip as proteins bind to the antibody-functionalized surface. In the resulting ratiometric video, individual proteins appear as transient signals upon landing, manifesting as localized dark pointspread functions, owing to destructive interference, before fading into the static background upon immobilization. This temporal isolation is crucial for particle tracking, enabling the algorithm to pinpoint and characterize unique binding events at the precise moment of adsorption, without interference from previously bound molecules. Ratiometric processing uses a sliding-window approach with a bin size of *N* = 150 frames, defined as the number of frames averaged in each of the two adjacent windows (one for the pre-event denominator and one for the post-event numerator) at a 500 Hz acquisition rate. The ratiometric frame at time step *k* (for pixel (*x, y*)) is given by

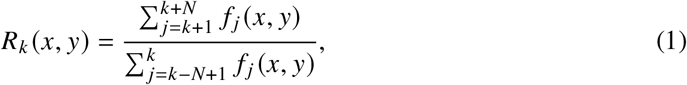

where *f* _*j*_ (*x, y*) are the high-pass filtered frames, yielding values centred around 1 in background regions. Plotting the contrast at a binding pixel over successive *k* yields a characteristic V-shaped profile: a linear downward slope starting at 1 down to the maximum destructive contrast (when the denominator contains no particle signal and the numerator fully includes it), followed by a linear upward slope back to 1 as the immobilized protein enters the background. This apex precisely marks the landing frame. This binning is essential to achieve an adequate signal-to-noise ratio (SNR) for quantifying protein contrast, as averaging multiple frames suppresses shot noise and other fluctuations that would otherwise obscure the weak scattering signals. The transient protein signal appears as a deviation from this baseline, so the ratiometric contrast is expressed as

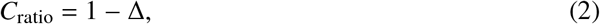

where Δ is the small positive fractional perturbation induced by the bound protein (on the order of 10^−3^ to 10^−2^ for proteins in the 100–1000 kDa range), and Δ scales linearly with the protein’s molecular mass (contrast deviation Δ ∝ mass, calibrated empirically for the biomolecular class). Such processing is indispensable given that protein scattering induces less than a 1% change in image contrast, which remains largely imperceptible in raw frames amid coverslip roughness and other noise, thereby enabling high-throughput, mass-resolved detection of single-molecule immuno-recognition in real time.

### Particle detection and contrast measurement

The You Only Look Once (YOLO v8) object detection model detects single landing events. The model was trained using a database of 2023 annotated ratiometric images from iSCAT experiments. Particles are uniquely labelled with an ID number and collected in a list. The Python script saves a 30 × 30 pixel image of the particle for each frame in which it appears, along with its location within the field of view. The data processing routine fits a two-dimensional Gaussian function to the ratiometric image of every detected particle. Peak amplitudes of fitted functions determine the particle contrasts in each frame.

The background noise for each experiment is quantified by computing the standard deviation of pixel intensity values within 20 selected 30 × 30 pixel squares at pseudo-random *xy* positions in each frame, then selecting the lowest value.

### Enzyme-linked immunosorbent assay

A Human IgM ELISA Kit (ThermoFisher Scientific, Cat: 88-50620-22) was used as directed. The IgM standard was tested in duplicate, and the human serum was tested in triplicate with a dilution of 10,000×. A Microplate Reader Multi-Mode FilterMax F5 measured the absorbance profile at 450 nm.

## Author Information

### Authors and Affiliations

**Department of Chemistry, University of British Columbia, Vancouver, BC V6T 1Z1 Canada** Carraugh C. Brouwer, Flaminia M. Muratori, Luke Melo, Edward Grant

**Biomedical and Materials Engineering, Politecnico di Milano, 20133 Italy** Flaminia M. Muratori

**Institute for Molecular and Translational Cardiology (IMTC), IRCCS Policlinico San Donato, Milan, 20097 Italy**

Flaminia M. Muratori, Luigi Anastasia, Carlo Pappone

**School of Medicine, University Vita-Salute San Raffaele, Milan, 20132 Italy**

Luigi Anastasia, Carlo Pappone

## Contributions

EG and LA conceived the experiment. CB and FM developed measurement protocols, prepared samples and substrates, gathered ratiometric images, and processed data. LM designed and built the interferometric microscope. LA and CP directed immunosorbent assays in Milan. All authors contributed to the writing and revision of the manuscript. EG and CP acquired the funding. EG supervised the project.

## Data and materials availability

All data needed to evaluate the conclusions in the paper are present in the paper and/or the Supplementary Materials. Raw data supporting this research are available via the following link: https://github.com/grantlab-ubc/immunoassay-iscat

