## Supplementary Materials for "Real-time mass-resolved label-free single-molecule immunoassay"

- Interferometric Scattering Contrast
- iSCAT instrumentation
- Micrograph videos
- Negative controls
- iSCAT landing rates on immunoassay coverslip as a function of IgM concentration
- Numbers of landings in 30-second measurements as a function of IgM concentration
- Standard addition of IgM to human serum
- iSCAT contrast as a function of molecular mass
- Enzyme-linked immunosorbent assay (ELISA)

### Interferometric Scattering Contrast

Interferometric scattering (iSCAT) microscopy detects a nanoparticle by measuring the interference between the light it scatters and a much stronger reference field reflected by an adjacent interface (*1, 2*). The detected optical intensity varies in proportion to the squared magnitude of the coherent sum of these two fields,

$$I_{\text{det}} = |E_r + E_s|^2 , \tag{1}$$

where $E_r$ is the reference (reflected) field and $E_s$ is the field scattered by the particle.

We express both fields in terms of the incident field amplitude $E_i$,

$$E_r = rE_i, \qquad E_s = sE_i e^{i\Delta\phi}, \tag{2}$$

where $r$ and $s$ are the (real) amplitude reflection and scattering coefficients, respectively, and $\Delta\phi$ is the phase difference between the scattered and reflected fields at the detector. The detected intensity then becomes

$$I_{\text{det}} = E_i^2 \left( r^2 + s^2 + 2rs\cos\Delta\phi \right) . \tag{3}$$

In the absence of a scattering particle, the detected signal arises purely from reflection at the interface,

$$I_{\text{bg}} = |E_r|^2 = E_i^2 r^2, \tag{4}$$

which defines the optical background. The signal attributable to the particle manifests in the intensity difference relative to this background,

$$\Delta I \equiv I_{\text{det}} - I_{\text{bg}} = E_i^2 \left( s^2 + 2rs\cos\Delta\phi \right) . \tag{5}$$

A conventional iSCAT experiment records a *contrast* defined by the particle scattering signal normalized to the background intensity (*3*),

$$C \equiv \frac{\Delta I}{I_{\text{bg}}} = \frac{s^2}{r^2} + \frac{2s}{r}\cos\Delta\phi. \tag{6}$$

When detecting weak scatterers such as proteins, the reflected field overwhelms the pure scattering field, such that

$$2rs \gg s^2. \tag{7}$$

Physically, this corresponds to the regime in which the interference (heterodyne) term dominates the detection, as opposed to a directly measured scattered intensity

$$I_{\text{det}} = E_i^2[r^2 + 2rscos\Delta\phi] \tag{8}$$

In this limit, the quadratic scattering contribution can be neglected, and the contrast reduces to the linearized expression

$$C \approx \frac{2s}{r} \cos \Delta\phi. \tag{9}$$

This linear dependence of contrast on the scattering amplitude $s$ is a defining feature of iSCAT microscopy. The explicit dependence on $\cos \Delta\phi$ points to the intrinsic phase sensitivity of the iSCAT, maximized by focus control.

Ratiometric signal processing captures a record of transient scattering signals against a dynamic background (*4*).

$$\frac{I_{\text{det}}}{I_{\text{bg}}} = 1 + C \approx 1 + \frac{2s}{r} \cos \Delta\phi, \tag{10}$$

where the constant term represents the dominant reflected background light, which is continually refreshed with the addition of previously bound particles.

In the Rayleigh regime, where particle diameter is on the order of 10% of the illumination wavelength, the field scattered from particles is proportional to the polarizability of the particle, $\alpha$. The polarizability of a spherical particle can be approximated as

$$\alpha = 3\epsilon_m V \frac{\epsilon_p - \epsilon_m}{\epsilon_p + 2\epsilon_m}$$

where $V$ is the particle volume, and $\epsilon_p$, $\epsilon_m$ are the complex dielectric functions of the particle and medium, respectively. (*5*) When working with particles of similar chemical composition in the same medium, the dielectric functions contribute little to the variation in polarizability compared with the effect of particle volume. Because $s$ scales with the particle polarizability, and thus approximately with molecular mass for small biomolecules, interferometric amplification enables the mass-resolved detection of individual proteins far below the sensitivity limit of conventional scattering measurements.

### iSCAT instrumentation

Figure S1 diagrams the optical setup used for the current suite of iSCAT measurements. In the illumination path, light, emitted by a 520 nm broadband laser (LaserTack LDM-520-1500-C), passes through a half-wave plate (HWP: Thorlabs WPH05M-532), followed by a pair of acousto-optic beam deflectors (AOBDs: Gooch & Housego R45100-5-6.5DEG-51-X/Y, MLV050-90-2AC-A1). The AOBDs are modulated by a function generator (Tektronic AFG-1062), steering the laser beam in both the $x$ and $y$ directions, to form a square illumination field. This beam proceeds through a $4f$ lens system (L1, L2: $f$ = 200 mm), a polarized beam splitter (Thorlabs PBS251), a quarter wave plate (QWP: Thorlabs WPQ10M-532) to reach the backplane of a high numerical aperture oil-immersion objective (Olympus UPLAPO100XOHR, NA = 1.5, 100× magnification).

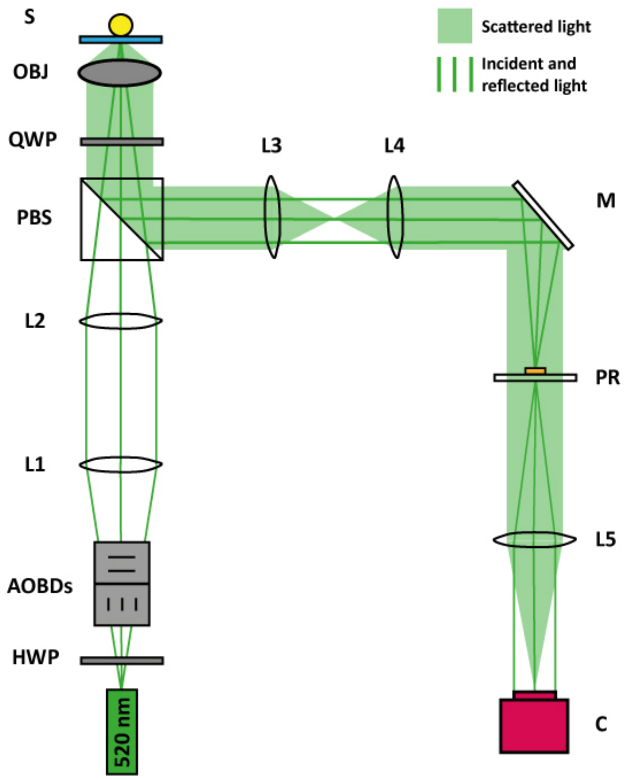


**Figure S1**: Schematic representation of the custom-built iSCAT instrument. The components present in the optical setup are a 520 nm diode laser, a half wave plate (HWP), acousto-optic beam deflectors (AOBDs), four lenses of focal length 200 mm (L1, L2, L3, L4), a polarized beam splitter (PBS), a quarter wave plate (QWP), a 100× objective (OBJ), a mirror (M), a partially reflective mask (PR), a lens of focal length 400 mm (L5), and the CMOS camera (C)

A custom-built sample stage holds a glass coverslip, optically coupled to the inverted objective by a drop of index-matching oil. Adjustment of a manual $z$-stage (Newport SDS65) provides rough focus. A piezoelectric actuator then moves the stage in the $z$-direction with nm precision, while a mechanical stage (OWIS KT 90 XY) enables manual movement along the $x$- and $y$-directions.

Reflected and scattered light generated at the coverslip-buffer interface returns along the illumination path and passes again through the QWP. This rotates the polarization by 90°, causing the polarizing beam splitter to steer the signal into the detection optical train. A second $4f$ lens system (L3, L4: $f$ = 200 mm) spatially separates the reflected and scattered beams. A partially reflective mask composed of a 3 mm-diameter, 150 nm-thick silver film at the centre of an NBK7 glass window attenuates the reflected light component at the conjugate focal plane of the AOBDs (*4*). Light crosses the last lens (L5, $f$ = 400 mm) to focus on the detection plane of a CMOS Blackfly camera (FLIR, Cat: BFS-U3-17S7M-C). Custom LabVIEW software provides a user interface for controlling the instrument.

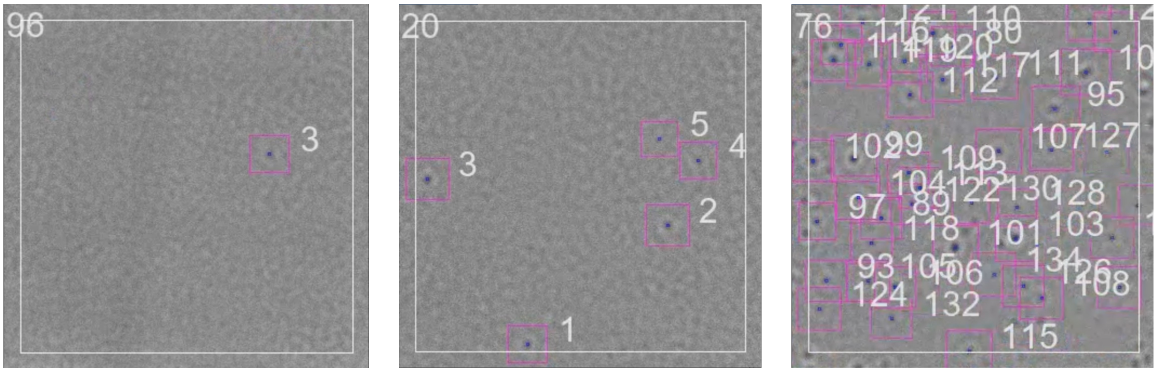


IgM 0.046 nM IgM 0.46 nM IgM 4.6 nM

**Figure S2**: Single frames taken from video records of interferometric signals produced by single-molecule IgM immunosorption on anti-IgM derivitized coverslips with casein blocking. These stills reflect the representative density of landings in a 0.4-second interval. IgM molecules appear as dark spots on a gray background before disappearing into the background. The image processing software creates videos from CMOS camera images collected at 500 frames per second, then temporally averaging with a bin size of 10 to produce a stream with an effective frame rate of 50 Hz. Our single-particle tracking algorithm identifies every landing, labelled on-screen with a box and a unique particle number. The number in the upper-left corner identifies the frame. These images show a field of view of 13.4$\mu$m × 13.4$\mu$m

### Micrograph videos

Ratiometric image processing, as described in the text, produces evanescent images directly signalling each event in which an IgM pentamer binds to an anti-IgM molecule affixed to the derivatized and blocked immunoassay coverslip. Figure S2 shows frames taken from experiments measuring landing rates for IgM solutions with concentrations of 0.046 nM, 0.46 nM, and 4.6 nM IgM in PBS.

The URL below links to short videos from which these screenshots were taken, showing the first 200 frames of iSCAT ratiometric videos of experiments observing binding events on immunoassay-prepared coverslips from 0.046 nM, 0.46 nM, and 4.6 nM IgM solutions in PBS.

https://github.com/grantlab-ubc/immunoassay-iscat/tree/main/IgM%20immunoassay%20videos

### Negative controls

We conducted negative-control experiments using PBS buffer to test whether the deposited casein and antibody particles caused particle-detection events (signal above background). The raw camera background was noticeably different for coverslips with deposited casein and antibody compared with PDL-only coverslips. Few landings were observed on coverslips with PBS buffer, with deposited casein and antibody blocked with casein. Overall, IgM and IgA detection proceeds in a background with a very low rate of false positives.

**Table S1**: Average landing rates observed in negative control experiments.

| Coverslip functionalization | Sample | Observed landing rate ($s^{-1}$) |
|---|---|---|
| casein | PBS buffer | 0.01 ± 0.002 |
| anti-Ig and casein | PBS buffer | 0.05 ± 0.04 |
| casein | IgM (5 nM) | 0.3 ± 0.2 |
| casein | normal serum (5000 × dilution) | 3.3 ± 0.6 |
| casein | depleted serum (5000 × dilution) | 1.5 ± 0.4 |
| anti-Ig and casein | ferritin (5 nM) | 0.06 ± 0.06 |

Negative controls using casein-functionalized coverslips assessed the effectiveness of casein blocking. Here, experiments tested landing rates for high-concentration IgM, normal serum, and depleted serum samples on casein-functionalized coverslips. For the IgM sample, casein blocked 99.5% of landings compared to IgM on PDL. The normal and depleted serum samples showed similar levels of suppressed landing frequency.

A highly concentrated sample of ferritin tested the discriminating power of the anti-Ig functionalized coverslip. Few landings were observed, establishing that the single-molecule immunoassay procedure has low affinity for this non-specific protein.

### Tabulated iSCAT landing rates of IgM on an anti-Ig functionalized immunoassay coverslip as a function of IgM concentration

Table S2 gives the accumulated number of landings as a function of time, sampled at one-second intervals for IgM pentamer mass-resolved landings on immunosorbent cover slips from solutions of the indicated IgM pentamer concentrations in PBS buffer. These data support the results in the main text displayed in Figure 1C.

**Table S2**: Cumulative number of IgM landing events as a function of time on an anti-Ig functionalized immunoassay coverslip sampled at one-second intervals in 30-second measurements for the indicated IgM concentrations in PBS buffer.

| time (s) | 0.046 nM | 0.093 nM | 0.46 nM | 0.93 nM | 2.32 nM | 4.64 nM |
|---|---|---|---|---|---|---|
| 0 | 0 | 0 | 0 | 0 | 0 | 0 |
| 1 | 0 | 1 | 7 | 21 | 59 | 47 |
| 2 | 0 | 2 | 16 | 45 | 107 | 120 |
| 3 | 0 | 6 | 19 | 63 | 155 | 193 |
| 4 | 0 | 9 | 23 | 83 | 203 | 276 |
| 5 | 1 | 9 | 34 | 97 | 247 | 319 |
| 6 | 2 | 12 | 43 | 111 | 294 | 395 |
| 7 | 2 | 17 | 47 | 134 | 348 | 465 |
| 8 | 2 | 19 | 53 | 148 | 375 | 530 |
| 9 | 4 | 20 | 59 | 163 | 408 | 591 |

*Continued on next page*

*Continued from previous page*

| time (s) | 0.046 nM | 0.093 nM | 0.46 nM | 0.93 nM | 2.32 nM | 4.64 nM |
|---|---|---|---|---|---|---|
| 10 | 4 | 22 | 66 | 182 | 452 | 664 |
| 11 | 4 | 24 | 71 | 195 | 486 | 738 |
| 12 | 4 | 26 | 73 | 217 | 521 | 818 |
| 13 | 5 | 29 | 76 | 230 | 571 | 888 |
| 14 | 5 | 32 | 81 | 248 | 607 | 954 |
| 15 | 6 | 32 | 88 | 260 | 637 | 1027 |
| 16 | 6 | 33 | 93 | 271 | 682 | 1098 |
| 17 | 7 | 35 | 95 | 287 | 714 | 1154 |
| 18 | 7 | 38 | 103 | 296 | 756 | 1223 |
| 19 | 7 | 41 | 107 | 306 | 790 | 1272 |
| 20 | 9 | 41 | 109 | 319 | 830 | 1327 |
| 21 | 9 | 41 | 113 | 330 | 870 | 1385 |
| 22 | 9 | 43 | 120 | 340 | 910 | 1454 |
| 23 | 9 | 44 | 128 | 348 | 946 | 1511 |
| 24 | 9 | 46 | 130 | 368 | 990 | 1579 |
| 25 | 9 | 50 | 133 | 375 | 1024 | 1641 |
| 26 | 9 | 51 | 137 | 387 | 1062 | 1688 |
| 27 | 11 | 54 | 144 | 402 | 1093 | 1743 |
| 28 | 11 | 56 | 148 | 408 | 1128 | 1804 |
| 29 | 12 | 57 | 152 | 419 | 1161 | 1859 |
| 30 | 12 | 60 | 157 | 430 | 1189 | 1914 |

Table S3 lists the number of IgM mass-resolved landings recorded in 30-second measurements as a function of IgM concentration in PBS buffer. These data support the results in the main text displayed in Figure 1D.

**Table S3**: Total number of IgM landing events on an anti-Ig functionalized immunoassay coverslip detected in 30-second measurements for the indicated IgM concentrations in PBS buffer.

| IgM concentration (nM) | Average number of detected landings | Standard error |
|---|---|---|
| 0.046 | 5.2 | 1 |
| 0.093 | 28 | 11 |
| 0.46 | 168 | 33 |
| 0.93 | 312 | 56 |
| 2.32 | 898 | 103 |
| 4.64 | 1643 | 162 |

### Standard addition of IgM to human serum

Table S4 lists the number of IgM mass-resolved iSCAT landings on immunosorbent coverslips recorded in 30-second measurements as a function of the added IgM concentration in a human serum sample. These data pertain to the results in the main text, plotted in Figure 2D, supporting the determination of native IgM in a human serum sample by the method of standard addition.

**Table S4**: Total number of IgM landing events on an anti-Ig functionalized immunoassay coverslip detected in 30-second measurements for the indicated IgM concentrations in a sample of human serum (5000 $\times$ dilution).

| Added IgM concentration (nM) | Average number of detected landings | Standard error |
|---|---|---|
| 0 | 83.5 | 16 |
| 0.5 | 289 | 33 |
| 1 | 392 | 126 |
| 2 | 1045 | 9 |
| 4 | 1530 | 37 |

### Variation of iSCAT contrast with protein mass

We optimized an iSCAT optical train with a reduced contrast range and sensitivity to focus on the point-spread functions of the IgM decamer, IgM pentamer, and IgA dimer. Calibration with a selection of additional standards establishes a precise linear relationship between iSCAT contrast and protein mass. Table S5 tabulates the means of the contrast histograms obtained in iSCAT measurements for the indicated protein complexes. Figure S3 plots the variation of the observed contrast histogram means with molecular weight for complexes of IgM, IgA, Cas9-GFP and ferritin.

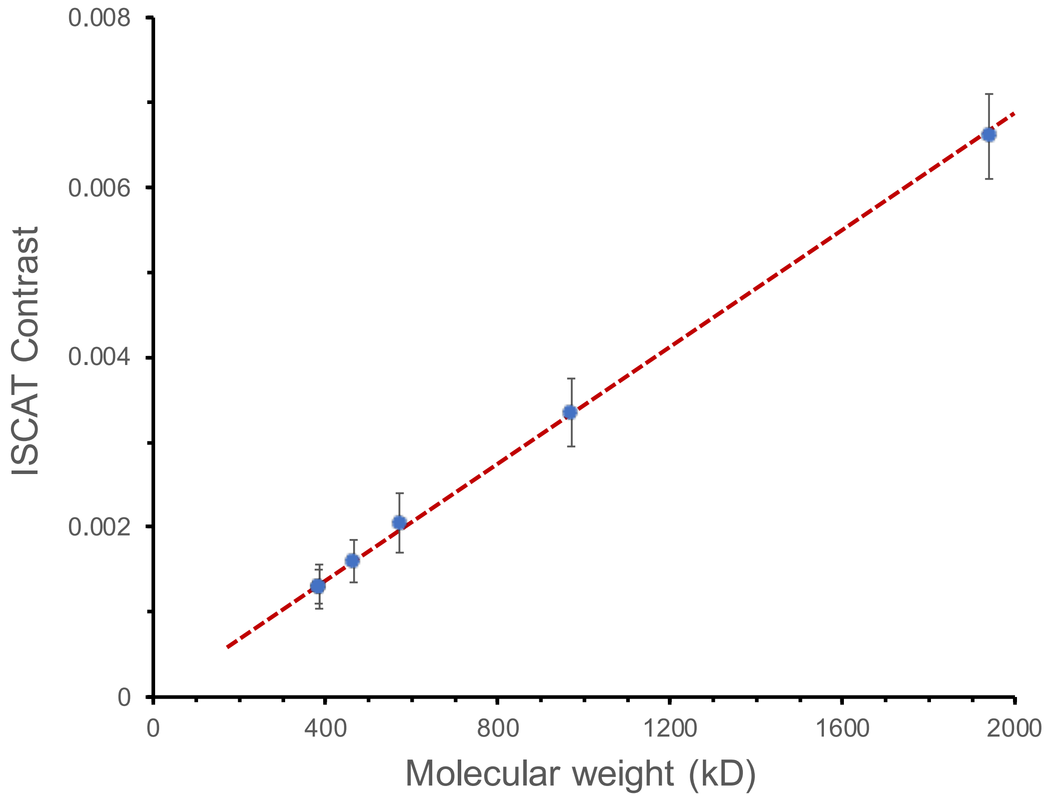


**Figure S3**: Molecular weight vs iSCAT contrast for proteins of known mass (IgA dimer, IgM pentamer & decamer, Cas9 GFP dimer & trimer, ferritin monomer).

### Enzyme-linked immunosorbent assay (ELISA)

Conventional measurements of specified proteins in biological samples often rely on the method of enzyme-linked immunosorbent assay (ELISA). Here, as a yardstick for the iSCAT analysis of IgM samples by direct counting of single-molecule binding events, we perform a conventional

**Table S5**: Observed iSCAT contrast as a function of biomolecule mass over a range from 384 to 1940 kDa.

| Protein | Molecular weight (kDa) | iSCAT contrast |
|---|---|---|
| IgM decamer | 1940 | 0.0066 |
| IgM pentamer | 970 | 0.0033 |
| Cas9-GFP trimer | 572 | 0.0020 |
| ferritin | 465 | 0.0015 |
| IgA dimer | 385 | 0.0013 |
| Cas9-GFP dimer | 384 | 0.0013 |

ELISA-based standard-addition experiment. After adding known amounts of IgM to a PBS buffer blank, we use ELISA to determine the analyte concentration. The results, listed in Table S6 and plotted in main text Figure 1E, deviate significantly from those expected of a well-defined standard addition experiment: (i) The response to added IgM exhibits a pronounced curvature above 200 ng/mL. (ii) The standard-addition response intercepts the $y$-axis with a measured nonzero response for a PBS blank.

**Table S6**: Measured absorbance in ELISA measures of IgM concentration standards.

| [IgM] ng/mL | 450 nm absorbance | Standard deviation |
|---|---|---|
| 15.6 | 0.0715 | 0.0007 |
| 31.2 | 0.0914 | 0.0005 |
| 62.5 | 0.1286 | 0.0011 |
| 125 | 0.2008 | 0.0013 |
| 250 | 0.3141 | 0.0163 |
| 500 | 0.4922 | 0.0499 |
| 1000 | 0.8875 | 0.0481 |

The observed curvature is consistent with well-known characteristics of conventional enzyme-linked immunoassays, in which antibody-antigen binding and enzymatic signal amplification yield a nonlinear transfer function. This arises from finite surface binding capacity, saturation of capture or detection antibodies, and non-proportional enzymatic turnover at higher levels of binding (*6, 7*).

The nonzero absorbance intercept at zero added IgM is more problematic, particularly for analyte detection at lower concentrations. In a true blank, a standard addition curve should intercept the origin. ELISA, however, commonly yields a false background concentration that can arise from a combination of sources. Residual enzyme-linked detection antibodies, nonspecifically adsorbed conjugates, or incomplete washing can generate product from chromogenic substrates, producing a finite optical density that adds to measurements at all levels of addition. Microplate absorbance measurements can include contributions from plate material that are not strictly zeroed by blank subtraction. Because an ELISA readout scales with enzymatic amplification rather than molecular count, it magnifies any residue of nonspecifically bound enzyme labels, producing a sizeable absorbance signal.

Contrast these limitations of ELISA with the iSCAT standard-addition measurement performed in human serum (main text Figure 2 D). Here, we observe a robust linear relationship between the added IgM concentration and the number of detected landing events across the full range studied. The linear fit extrapolates to a positive intercept of 83 counts over a 30 s window, corresponding directly to a native IgM concentration of 163 ng/mL in the unspiked serum sample.

Unlike ELISA, this intercept is physically meaningful. In iSCAT microscopy, the measured observable is the direct count of individual IgM adsorption events on an antibody-functionalized surface. The signal, therefore, scales linearly with analyte flux to the surface, provided that binding sites are not saturated and that depletion remains negligible over the observation window. Because there is no enzymatic amplification, chromogenic chemistry, or nonlinear optical readout, there is no mechanism for generating a concentration-independent offset. The intercept in iSCAT standard-addition thus reflects the true endogenous analyte concentration, rather than an instrumental background.
